# Whole-genome phylogenomics and synteny resolve a single origin of body-plan asymmetry in flatfishes

**DOI:** 10.64898/2026.05.25.727411

**Authors:** Julian Gallego-Garcia, Delson Hays, Pakorn Tongboonkua, Jeremiah J. Minich, Leon Hilgers, Todd P. Michael, Michael Hiller, Chao Zhang, Guillermo Orti, Dahiana Arcila, Wayne Pfeiffer, Emanuell Duarte-Ribeiro, Siavash Mirarab, Ricardo Betancur-R.

**Affiliations:** Marine Biology Research Division, Scripps Institution of Oceanography, University of California San Diego, La Jolla, CA, USA; Department of Fishery Biology, Faculty of Fisheries, Kasetsart University, Bangkok, Thailand; The Plant Molecular and Cellular Biology Laboratory, The Salk Institute for Biological Studies, La Jolla, CA, USA; Department of Biology, Baylor University, Waco, TX, USA; Senckenberg Research Institute, Frankfurt, Germany and Institute of Cell Biology and Neuroscience, Faculty of Biosciences, Goethe University Frankfurt, Frankfurt, Germany; Department of Cell and Developmental Biology, School of Biological Sciences, University of California San Diego, La Jolla, CA, USA; Department of Science and Conservation, San Diego Botanical Garden, Encinitas, CA, USA; Center for Marine Biotechnology and Biomedicine, University of California San Diego, La Jolla, CA, USA; School of Life Sciences, Peking University, Beijing, PR, China; San Diego Supercomputer Center, University of California San Diego, La Jolla, CA, USA; Department of Biological Sciences, The George Washington University, Washington, DC, USA; Zoological Institute, Department of Environmental Sciences, University of Basel, Basel, Switzerland; Department of Electrical and Computer Engineering, University of California San Diego, La Jolla, CA, USA

**Keywords:** Pleuronectiformes, synteny-based phylogenomics, rapid radiation

## Abstract

Flatfishes display the most dramatic asymmetric body plan in vertebrates, yet whether this rare innovation evolved once (flatfish monophyly, FM) or multiple times (flatfish polyphyly, FP) has remained contentious. A recent genome-wide study supported FP by placing *Psettodes*, the earliest-diverging flatfish lineage, among symmetric relatives within Carangaria, the clade that also includes billfishes, jacks, mahi-mahi, and barracudas. Subsequent work traced this to base-composition artifacts and inadequate substitution modeling. Here we revisit the question using whole-genome phylogenomic and synteny data from 17 carangarian species spanning flatfishes and carangarian relatives. We contribute three new chromosome-level assemblies, including the first for *Psettodes*. Nucleotide-based coalescent analyses (e.g., ROADIES, CASTER) yield strong support for FM, with *Psettodes* sister to all other flatfishes. Microsynteny analyses built from conserved gene-order blocks corroborate this result: topology tests, cluster-profile counts, and rearrangement-based trees favor FM over two competing FP topologies. Macrosynteny, based on chromosome-scale rearrangements, yields a more mixed signal, with support for FM depending on the metric and taxon-sampling scheme. We interpret this scale-dependent pattern in the context of the explosive post-Cretaceous radiation of Carangaria. The short intervals between speciation events that characterize rapid radiations appear to have left sufficient signal in fine-grained microsyntenic rearrangements, while chromosome-scale rearrangements were too rare to consistently resolve these closely spaced splits. When integrated with evidence from conserved developmental mechanisms active during metamorphosis, the stage at which flatfish asymmetry first emerges, and from the exceptionally complete fossil record, our multi-scale genomic evidence supports a single evolutionary origin of flatfish asymmetry.

## Introduction

Flatfishes possess the most extreme departure from bilateral symmetry among vertebrates. During post-larval metamorphosis, one eye migrates across the skull midline to the opposite side, yielding an adult that lies permanently on one side with both eyes directed upward (Gibson, 2005). The result is a body plan so radical that it once challenged the plausibility of evolutionary gradualism itself: G. Mivart^1^ cited it as evidence against Darwinian selection, and R. Goldschmidt^2^ invoked it as a textbook case for saltation, arguing that no functional intermediate between a symmetric ancestor and a fully asymmetric flatfish could be envisioned. That argument was laid to rest nearly a century later when M. Friedman^3^ described two Eocene fossils (*Amphistium paradoxum* and *Heteronectes chaneti*) whose partially migrated orbits demonstrated that the transition proceeded through incremental steps. A second longstanding puzzle, the interrelationships of flatfishes within the broader fish tree, was substantially advanced by molecular phylogenetics, which unexpectedly placed flatfishes alongside morphologically disparate lineages such as billfishes, remoras, archerfishes, mahi-mahi, and jacks within a clade named Carangaria^4–9^. In the pre-DNA era, morphology alone did not anticipate this grouping, reflecting the paucity of clear anatomical synapomorphies linking flatfishes to any specific spiny-finned fish lineage^10,11^. But a third and now more consequential question remains contentious: did flatfish body-plan asymmetry evolve once, or independently more than once within carangarians?

The answer carries profound evolutionary implications. If flatfishes are monophyletic, the entire developmental program underlying body-plan asymmetry (thyroid-hormone-driven metamorphosis^10,11^, *bmp4*-mediated pseudomesial bar formation^12^, and coordinated changes in pigmentation, skeletal remodeling, and lateralized behavior) evolved a single time. If polyphyletic, vertebrate evolution independently assembled this radical morphological innovation at least twice. Molecular phylogenetic studies are split on this question: some resolve flatfish monophyly (FM) (e.g.,^4,9,12,13^), whereas others place *Psettodes*—the critical early-diverging flatfish lineage—closer to non-flatfish carangarians (flatfish polyphyly, FP), implying dual origins of asymmetry (e.g.,^9,14,15^). Morphological evidence from the fossil record and comparative anatomy broadly favors a single origin^3,16–18^, as do non-coding ultraconserved element (UCE) datasets analyzed under both homogeneous and nonhomogeneous substitution models^12,13,16^. By contrast, protein-coding datasets are prone to base-compositional nonstationarity (BCNS) artifacts that can mislead inference toward polyphyly^9,12^. This prompted an exchange in Nature Genetics between Lü et al.^15^, who reported polyphyly from genome-wide exon data, Duarte-Ribeiro et al.^12^ who recovered FM after accounting for base-compositional non-stationarity and gene-tree estimation error, and Lü et al.^19^ who maintained their original conclusion. The analytical drivers of this persistent discordance include heterotachy along flatfish branches, GC-content nonstationarity, and sensitivity to substitution-model choice^9,12^. Collectively, these issues make the flatfish problem one of the most contentious unresolved nodes in the vertebrate tree of life, and arguably among the more prominent unresolved questions in animal phylogeny as a whole, perhaps second only to the Ctenophora-versus-Porifera debate^20,21^. Even so, the question has received far less attention than its scope would suggest.

Every molecular study on flatfishes to date has relied exclusively on nucleotide or amino-acid sequence data. None has tested flatfish monophyly using genomic structural characters. Yet synteny-based phylogenomics has recently helped resolve other difficult nodes: conserved gene linkage has been used to evaluate early animal relationships^20^, chromosome-scale rearrangements alongside gene sequences have informed the deepest split among living teleosts^22^, and deeply conserved synteny has clarified early vertebrate and metazoan chromosome evolution^23,24^. Microsynteny-based phylogenies have also been applied successfully to angiosperms^25^. As rare genomic changes, gene-order rearrangements are expected to show low phylogenetic noise (homoplasy)^26^, making them an attractive independent marker class for short-internode problems where sequence-based signal is ambiguous. At the same time, whole-genome phylogenomic methods now enable analyses that move beyond pre-selected loci. For example, CASTER infers species trees from site patterns extracted from whole-genome alignments^27^, whereas ROADIES automates species-tree inference directly from genome assemblies without relying on annotation or orthology detection^28^. Despite these advances, neither whole-genome phylogenomics nor synteny-based approaches have been applied to the flatfish problem.

Here we combine whole-genome phylogenomics with multi-scale synteny to re-evaluate the origin of the flatfish asymmetry. We generated a chromosome-level genome for *Psettodes erumei*—a representative of the lineage whose placement toggles inference between FM and FP—using PacBio Revio and Hi-C, and assembled a 17-species comparative dataset for Carangaria, including two additional new assemblies. We test FM against competing FP alternatives using three independent lines of evidence: (i) genome-wide coalescent phylogenomics from assemblies and whole-genome alignments (ROADIES, CASTER), complemented by TOGA2 single-copy orthologs and gene-genealogy interrogation^27–30;^ (ii) microsynteny phylogenomics using network-based clustering of collinear blocks (syntenet) and topology-dependent ancestral gene-order reconstruction (AGORA)^31,32^; and (iii) macrosynteny analyses combining Oxford grids, fusion-with-mixing simulations, and ancestral karyotype reconstruction with DESCHRAMBLER^20,24,33^. Genome-wide sequence and microsynteny results converge on FM with strong support, whereas macrosynteny yields a more mixed but FM-leaning signal. When integrated with extensive evidence from the fossil record and shared conserved developmental mechanisms active during metamorphosis, including thyroid hormone and BMP pathways^12,34,35^, our results support a single evolutionary origin of the flatfish asymmetric body plan.

## Results

### Nucleotide-based whole-genome phylogenomics supports flatfish monophyly

We first evaluated the competing hypotheses using nucleotide-based phylogenomic approaches applied to whole-genome data (Fig. 1), including ROADIES^28^, a reference-free pipeline that identifies homologous regions directly from genome assemblies. ROADIES selected 4,208 single- and multi-copy loci after 2 iterations from all 17 species, placing *Psettodes erumei* as sister to all other flatfishes with maximum support (ASTRAL-Pro2 local posterior probability = 1.0), favoring FM. To incorporate information from whole-genome data, we performed two independent Progressive Cactus^36^ alignments (workflow in Supplementary Fig. S4) using guide trees that reflected the monophyly and polyphyly hypotheses, respectively, and extracted them under two reference projections (the outgroup *Mastacembelus armatus* and *P. erumei*). Because raw CASTER analyses of these alignments systematically recapitulated the input guide-tree topology, indicating propagation of guide-tree bias into species-tree inference, we developed a post-processing pipeline involving re-alignment with MAFFT and trimming with trimAl to reduce this dependence (see Methods). Post-processed alignments comprised over 120 million sites each, of which approximately 33 million were parsimony-informative. Three of four post-processed CASTER analyses resolved FM: both CASTER-site and CASTER-pair on the monophyly-guided alignment (100% bootstrap), and CASTER-site on the polyphyly-guided alignment (87.2% bootstrap). The single exception, CASTER-pair on the polyphyly-guided alignment, recovered an unexpected topology in which *Eleutheronema tetradactylum* (a non-flatfish carangarian) was placed as sister to Pleuronectoidei (99.9% bootstrap), with *Psettodes* sister to that clade implying flatfish paraphyly rather than polyphyly (Fig. S5). Robinson-Foulds comparisons confirmed that post-processed trees converged rather than simply recapitulating their input guide trees (Supplementary Figs. S9, S10).

**Figure 1.**
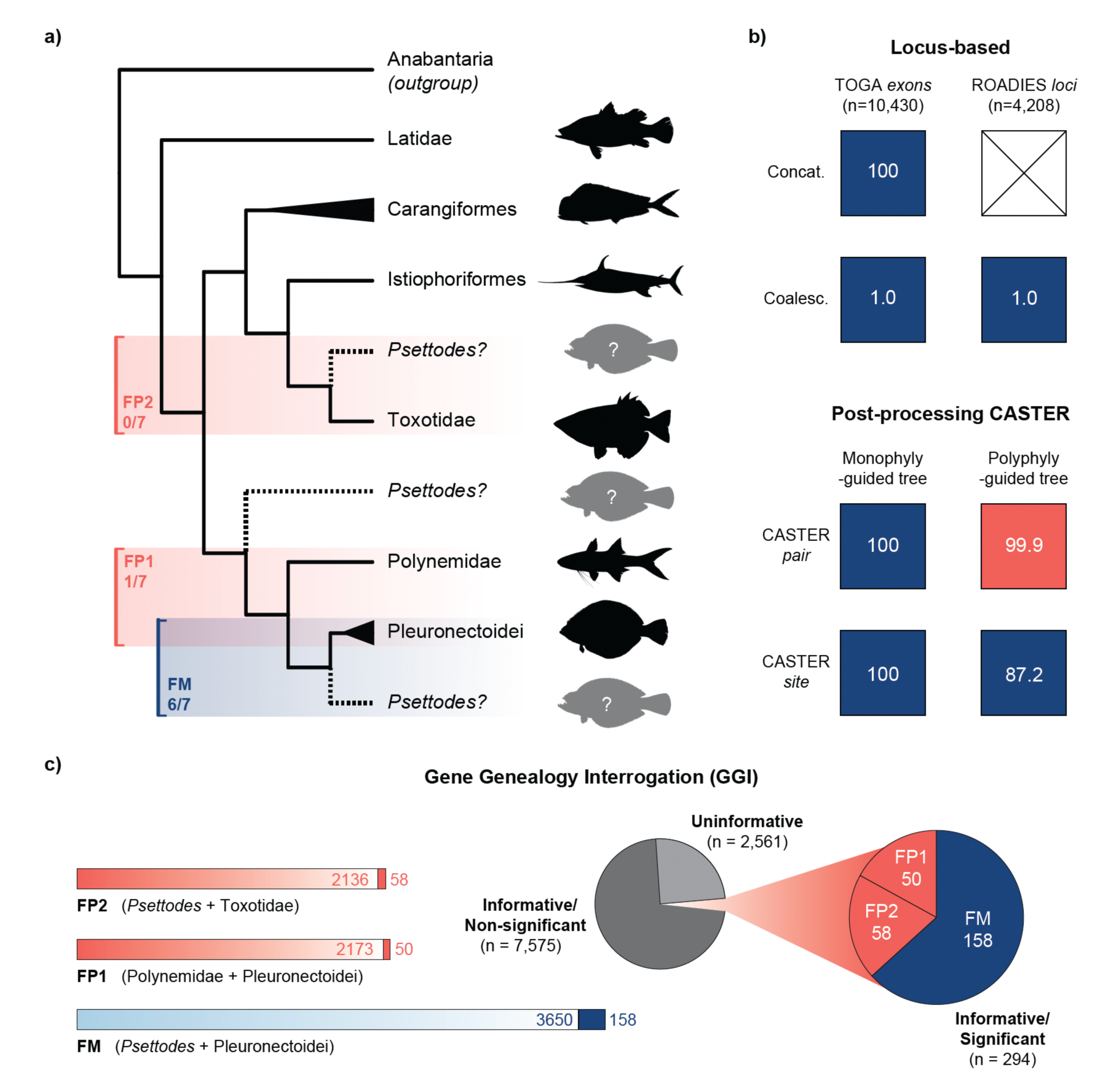
Summary of nucleotide-based phylogenomic support for flatfish monophyly. **A**, Schematic of three focal alternatives: flatfish monophyly (FM), with *P. erumei* sister to sampled pleuronectoids; flatfish polyphyly 1 (FP1), with *Eleutheronema tetradactylum* (Polynemidae) sister to sampled pleuronectoids; and flatfish polyphyly 2 (FP2), with *Psettodes erumei* sister to *Toxotes jaculatrix* (Toxotidae). **B**, Matrix summarizing results from ROADIES, a TOGA2 ASTRAL analysis, and post-processed monophyly- and polyphyly-guided CASTER analyses (for pre- processing CASTER analyses see Figs. S5–S8). Number insets indicate node support for the resultant topology (bootstrap for IQ-TREE and CASTER; local posterior probability for ASTRAL/ROADIES). **C**, gene- genealogy-interrogation (GGI) results based on the AU topology test, showing the number of genes non-significantly (light bars) and significantly (dark bars) supporting each node.

Independent validation came from 10,430 exon trees based on high-quality one-to-one orthologs identified by TOGA2, from which ASTRAL-III resolved FM with maximum support (local posterior probability = 1.0; gene concordance factor = 21.9%; site concordance factor = 35.2%). A concatenation-based maximum-likelihood analysis of these loci, using a partitioning scheme in which individual genes were partitioned by codon position, also recovered flatfish monophyly with full support (bootstrap = 100). To further evaluate relative support at individual loci, we applied a gene-genealogy-interrogation (GGI) approach^30^ in which three constrained topologies were evaluated at every locus using AU topology tests: flatfish monophyly (FM: *Psettodes* + Pleuronectoidei), and two alternative polyphyly topologies, FP_1_ (*Eleutheronema* + Pleuronectoidei) and FP_2_ (*Psettodes* + *Toxotes*), both concordant with the tree favored by Lü et al.^15^. FM was rejected by the smallest fraction of genes (5.9%), compared with 10.6% for FP_1_ and 8.7% for FP_2._ Among the 294 strongly discriminative loci (p-AU ≥ 0.95 for the top-ranked topology), 186 (63.3%) favored FM, 58 (19.7%) favored FP_1_, and 50 (17.0%) favored FP_2_. Collectively, 10 of 11 nucleotide-based analyses supported FM (Fig. 1).

### Microsynteny phylogenomics resolves *Psettodes* as sister to Pleuronectoidei

We next asked whether conserved local gene order provided independent support for FM, using two complementary microsynteny frameworks: syntenet, which clusters protein-based collinear blocks into a binary character matrix, and AGORA, which reconstructs ancestral gene-order relationships from TOGA2-derived single-copy orthologs. All results are summarized in Figure 2.

**Figure 2.**
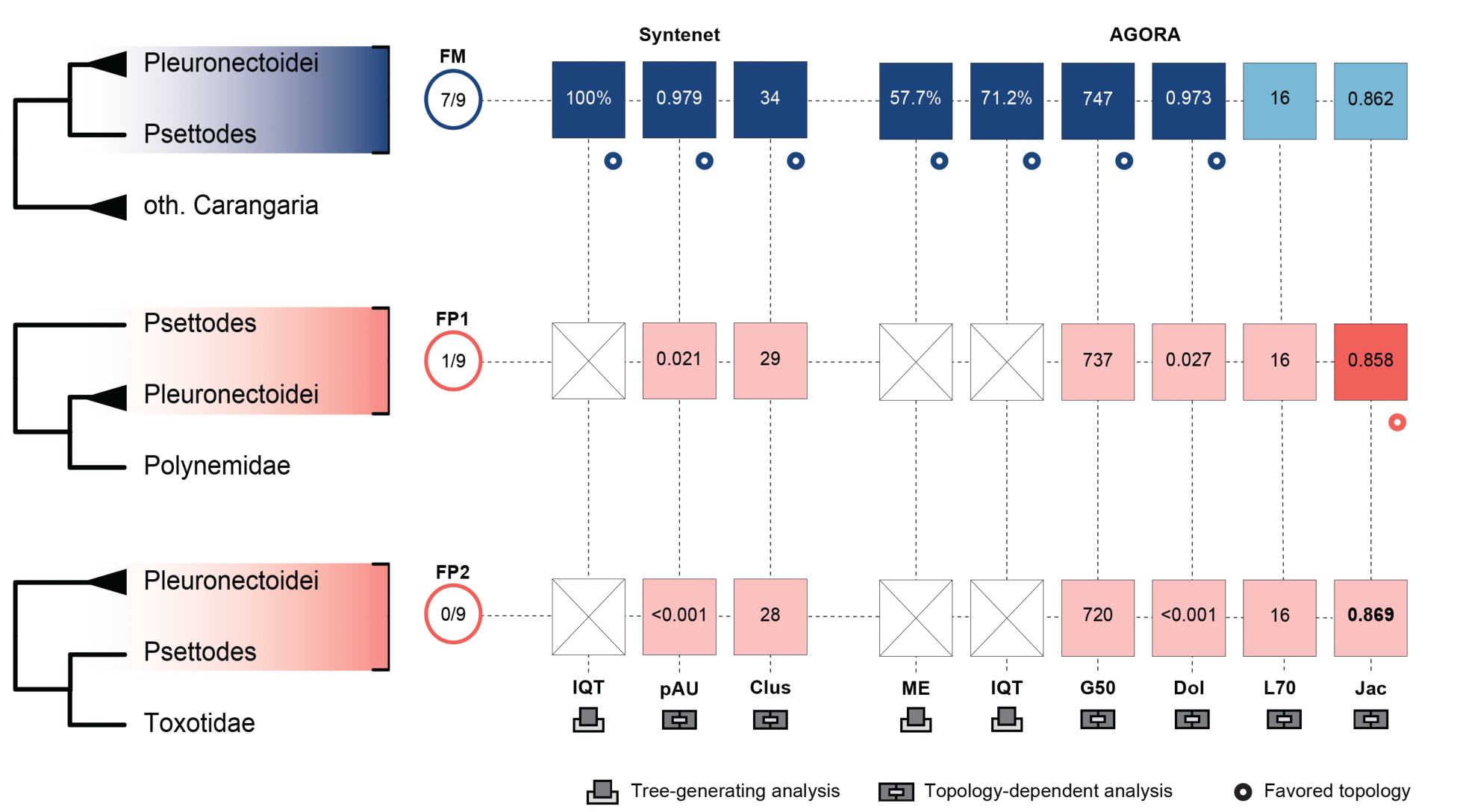
Summary of microsyntenty-based support for three topological hypotheses (FM, FP1, FP2). Rows correspond to nodes FM, FP1, FP2. Columns correspond to ten tests using output of either syntenet or AGORA, divided into tree-generating (n = 3) and tree-dependent (n = 7) tests: IQT (IQ-TREE ultrafast bootstrap support), ME (FastME tree bootstrap support), pAU (approximately-unbiased topology test weight), Clus (number of topology-exclusive clusters), G50 (AGORA ancestral-region contiguity statistic), Dol (Akaike weight from the Dollo-likelihood adjacency model), L70 (AGORA ancestral-region contiguity statistic), and Jac (Jaccard similarity to the extant *P. erumei* adjacency set). Number insets indicate quantitative support for each topology per metric. Per-metric values are reported in SM Tables S12–S16; corresponding visualizations appear in SM Figs. S18–S25.

The syntenet analysis produced 23,850 binary microsynteny characters for 17 taxa that were retained after DIAMOND BLASTP filtering at >70% amino-acid identity and >30 aa aligned length, followed by MCScanX collinearity detection requiring at least five anchors per block and a maximum gap of 25 genes (workflow in Supplementary Fig. S15; pairwise microsynteny-conservation heatmap in Supplementary Fig. S16). A maximum-likelihood tree inferred under the Mk model using the resulting matrix resolved FM with 100% ultrafast bootstrap support (Supplementary Fig. S18). When the same matrix was used to compare the three candidate topologies via AU tests (Supplementary Fig. S19), FM provided the best fit (p-AU = 0.979), FP_1_ was rejected (p-AU = 0.021), and FP_2_ was strongly rejected (p-AU = 8.97 × 10⁻⁹). Of the 23,850 microsynteny clusters, 20,579 were retained after excluding singletons and those present in more than 90% of taxa. Of this filtered set, topology-exclusive clusters (present in every species united by a candidate topology and absent from the alternatives) totaled 34 for FM, 29 for FP_1_, and 28 for FP_2_ (Supplementary Table S13).

The AGORA gene-order analyses reinforced these results. A rearrangement-distance minimum-evolution tree (FastME) resolved FM as sister to Pleuronectoidei with 57.7% bootstrap support (Supplementary Figs. S20, S21; sensitivity analysis in Supplementary Figs. S22, S23), and the topology was more similar to the FM constraint than to either alternative (normalized RF = 0.250 vs 0.333). A maximum-likelihood tree inferred from a binary gene-adjacency matrix likewise resolved FM (71.2% SH-aLRT; Supplementary Fig. S24; Supplementary Table S14). In constrained three-topology AU tests on this adjacency matrix, FM had the highest likelihood (p-AU = 0.702), but neither alternative was statistically rejected (FP₁: p-AU = 0.347; FP₂: p-AU = 0.365). Ancestral gene-order reconstructions under each topology revealed a consistent pattern: extant adjacency parsimony favored FM in four AGORA configurations (a strict vertebrates run retaining orthologs at score ≥ 0.99; a relaxed generic run; a relaxed vertebrates run, our primary configuration; and a *Solea solea*-excluded relaxed vertebrates sensitivity run), with bootstrap unique-best fractions always strongest for FM (0.789, 0.955, 0.955, and 0.966, respectively). Ancestral contiguity (G50) also favored FM, and incident breakpoints were lowest under FM (G50 = 747 under FM versus 738 under FP₁ and 720 under FP₂; 1,813 incident breakpoints under FM versus 1,844 under FP₁ and 2,231 under FP₂). A Dollo-likelihood analysis (Supplementary Table S16) of derived gene adjacencies provided the best fit under FM (Akaike weight = 0.973), with FP₁ receiving negligible support (ΔAIC = 7.17; Akaike weight = 0.027) and FP₂ strongly disfavored (ΔAIC = 38.26). In total, across the AGORA ancestral-reconstruction metrics (Supplementary Table S15), the majority of comparisons favored FM, one favored FP₂, and none favored FP₁; the L70 contiguity comparison was non-discriminatory (L70 = 14 for FM, FP₁, and FP₂ in the strict-vertebrates run; L70 = 16 for all three in the relaxed-generic run; Table S5). We note that individual tree-generating synteny methods varied in their internal resolution, and some recovered locally weakly supported or biologically implausible nodes away from the focal *Psettodes* placement. The aggregate signal nevertheless converged on FM.

### Macrosynteny analyses are FM-leaning but inconclusive

To test whether chromosome-scale rearrangements carry phylogenetic signal for the three competing topologies, we combined multispecies and pairwise protein-based comparisons with DESCHRAMBLER ancestral chromosome reconstruction. A multispecies ribbon plot built from an expanded ancestral linkage group (ALG) homology table (Supplementary Figs. S28) summarized chromosome-scale structure across all 17 carangarian taxa, using 24 Carangaria-specific ALGs resolved by overlapping four-way reciprocal-best-hit analyses (Figure 4). Most non-flatfish carangarians retained largely one-to-one or simple fused correspondences with individual ALGs, with any additional large-scale rearrangement restricted to intrachromosomal inversions, whereas pleuronectoid genomes showed repeated fusion, fission, and mixing of ALG segments between chromosomes. *Eleutheronema tetradactylum* was the exception among non-flatfish carangarians, with its derived set of 26 chromosome-scale scaffolds (versus n = 24 in most carangarians) consistent with a lineage-specific, autapomorphic fission (Supplementary Fig. S27). This concentration of structural change in pleuronectoids is consistent with broader vertebrate-scale reconstructions in which the tongue sole (*Cynoglossus semilaevis*) stands out among the most extensively rearranged vertebrate genomes^32^ Descriptively, the ribbon-plot pattern contextualizes the focal hypotheses but does not by itself discriminate among them.

**Figure 3.**
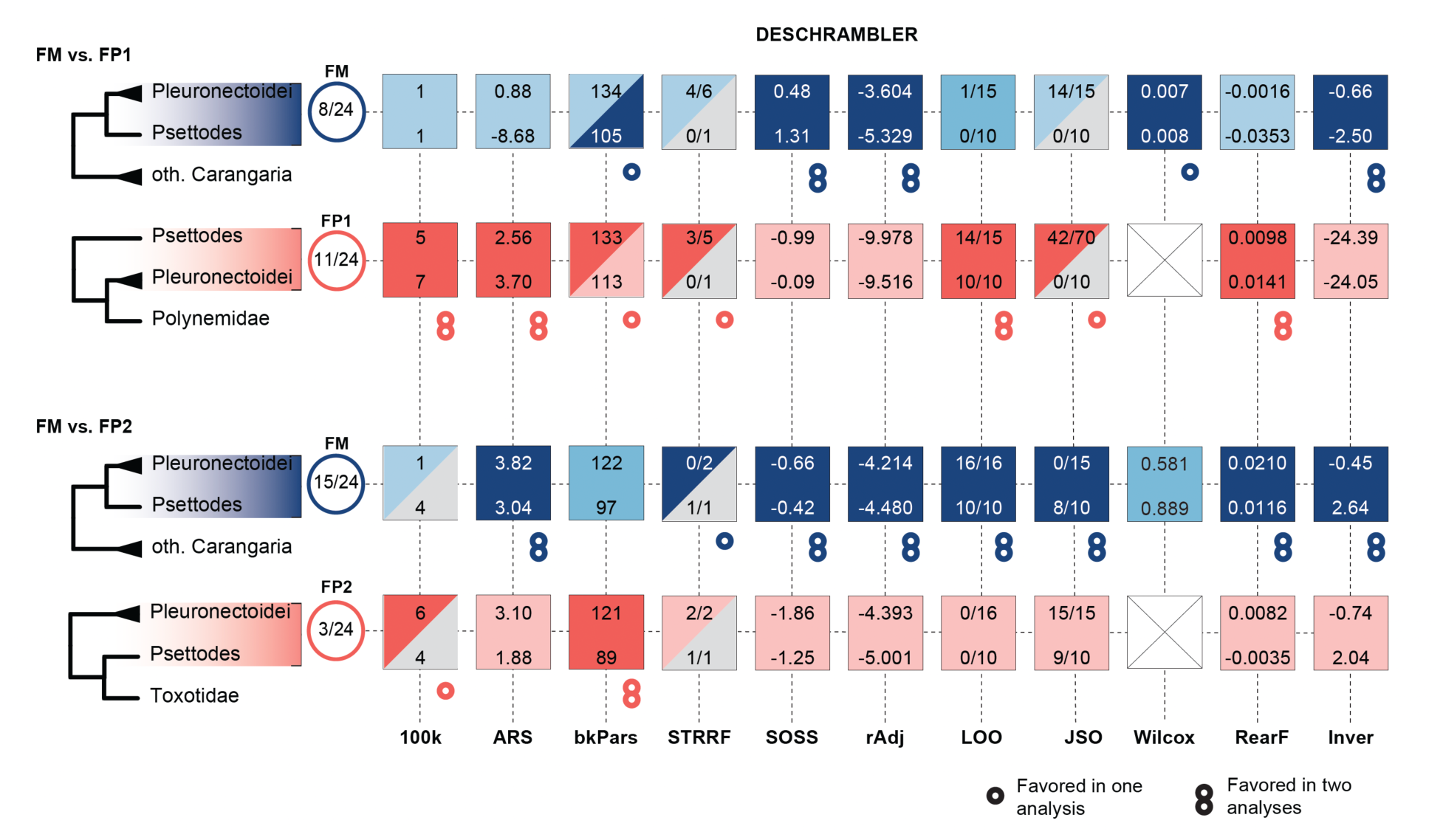
Summary of macrosyntenty-based support for two topological comparisons (FM vs FP1 and FM vs FP2). Columns correspond to twelve tree-dependent tests using output of DESCHRAMBLER: 100k (100-kb raw-metric score), ARS (adjacency-retention separation), bkPars (breakpoint parsimony), STRRF (switching-taxon retention role-fit), SOSS (switching-outgroup split support), rAdj (raw adjacency likelihood), LOO (leave-one-out separation wins), JSO (jackknife switched-outgroup descendant-like fraction), Wilcox (Wilcoxon paired-coordinate matched-adjacency test), RearF (per-APCF rearrangement fraction), and Inver (inversion-rate). Number insets indicate quantitative support for each topology per metric. Per-metric values are reported in SM Tables S21 and S23; corresponding evidence-matrix visualizations appear in SM Figs. S35–S38.

**Figure 4.**
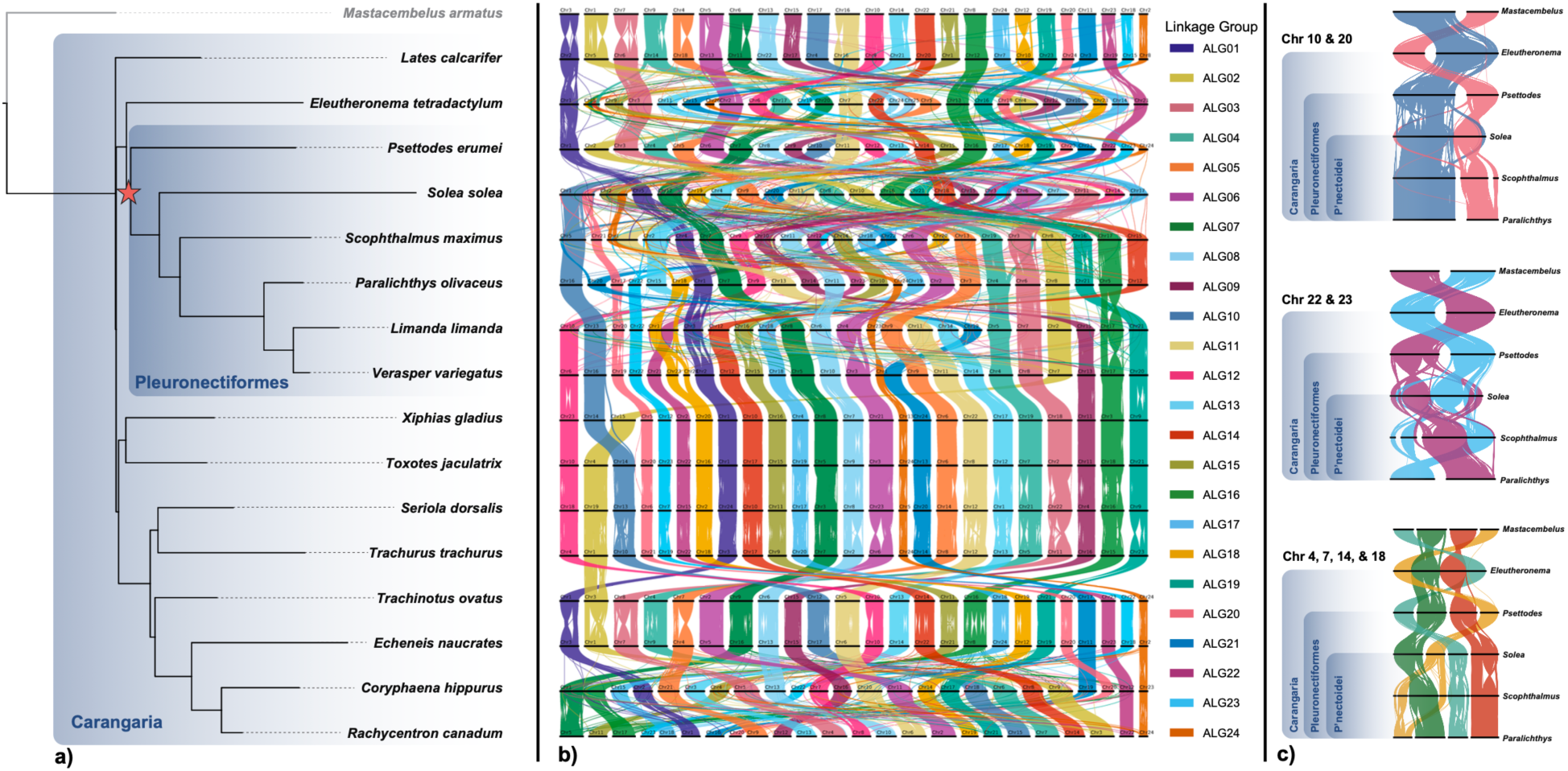
Macrosynteny ribbon plot. (a) ROADIES species tree for the 17 sampled carangarian genomes; the red star marks the flatfish-monophyly node. (b) Ribbon plot showing multispecies ancestral linkage group (ALG) correspondences (ALG01–ALG24) across the sampled genomes, drawn in a fixed taxon order to allow direct visual comparison of chromosome-scale structure. Panel (b) uses the FM topology (*Psettodes erumei* + Pleuronectoidei); the corresponding ribbon plots reconstructed under the alternative FP₁ and FP₂ topologies are shown in Figure S28. (c) Selected chromosome comparisons across representative carangarian genomes, highlighting focal chromosomes whose fusion/fission events and ALG-segment rearrangement or mixing are concentrated in pleuronectoid lineages.

The deeper Bilaterian-Cnidarian-Sponge^24^ and chordate^23^ linkage-group frameworks were consulted to distinguish conserved linkages from lineage-specific rearrangements. Interpreted against this backdrop, pairwise Oxford Dot Plot comparisons among the 17 carangarian species showed that chromosome-scale fusion, fission, and mixing were concentrated in Pleuronectoidei. Topology-dependent tests then asked whether any of these rearrangements carried signal diagnostic of FM, FP₁, or FP₂. Across multiple quartet and quintet configurations, fusion-with-mixing simulations did not provide decisive support for any topology (Supplementary Figs. S29–S33); the signal was concentrated in derived rearrangements in Pleuronectoidei, particularly in *Solea solea*, rather than in clear synapomorphies diagnostic of FM, FP₁, or FP₂ (Supplementary Fig. S27). Together, pairwise and multispecies protein-based comparisons leaned toward FM without rejecting FP₁ or FP₂, prompting a complementary ancestral-chromosome reconstruction.

DESCHRAMBLER ancestral chromosome reconstructions were evaluated in two reference-specific pairwise comparisons that share FM: a *Scophthalmus*-anchored FM versus FP1 comparison and a *Psettodes*-anchored FM versus FP_2_ comparison, each under full-taxon-sampling and more balanced two-descendant-only designs (Supplementary Fig. S34; Supplementary Tables S7, S8, S9). In all four comparisons, adjacency log-likelihood favored FM, with FM-alternative gaps ranging from 0.18 log-likelihood units (*Scophthalmus*-anchored, full-taxon: −4.214 for FM versus −4.393 for FP1) to 6.37 units (*Psettodes*-anchored, full-taxon: −3.604 for FM versus −9.978 for FP2). Formal same-reference tests mirrored this asymmetry: results were inconclusive under the *Scophthalmus* reference (APCF-cluster bootstrap 95% CI spanning zero; paired Wilcoxon p = 0.581) but favored FM under the *Psettodes* reference (bootstrap mean Δlog-likelihood = 6.374, 95% CI 4.585–7.964; paired Wilcoxon p = 0.007). Fragmentation-style metrics and adjacency-likelihood metrics repeatedly pointed in opposite directions, reflecting the inherent difficulty of ancestral chromosome reconstruction when pleuronectoid genomes are among the most rearranged in the dataset. Overall, macrosynteny evidence leaned toward FM but lacked the statistical resolution to discriminate among topologies (Fig. 3).

## Discussion

### Why sequence-based phylogenomics has struggled with this node

The flatfish monophyly problem is difficult for sequence-based methods because the relevant internode is short, the flanking branches are long, and flatfish lineages exhibit pronounced heterotachy and GC-content nonstationarity^9,12^. Protein-coding loci are especially vulnerable: GC-biased gene conversion generates compositional attraction between lineages with convergently high GC content, mimicking the phylogenetic signal expected under polyphyly^9,12^. Non-coding UCE datasets, which are less affected by these artifacts, consistently resolve FM under both homogeneous and nonhomogeneous models^12,13^, and the nonhomogeneous GHOST model resolves FM even from exonic data when applied to datasets small enough for its parameter space^12,37^. Our whole-genome analyses extend this pattern. CASTER and ROADIES, which operate on genome-scale site patterns and randomly sampled genomic segments respectively, converge on FM from datasets orders of magnitude larger than any previously analyzed for Carangaria (over 120 million alignment sites; 4,208 ROADIES loci; 10,430 TOGA2 one-to-one orthologs). The consistency of FM recovery across these disparate methods and data sources indicates that the polyphyletic signal in earlier studies reflects systematic model misspecification rather than genuine phylogenetic discordance.

Our results also bear directly on the arguments advanced by Lü et al. in their reply^19^. First, their gene-wise ΔGLS analysis, which found that a majority of gene trees favored FP, relied exclusively on homogeneous substitution models applied to exonic data—the precise analytical combination shown to be most susceptible to BCNS artifacts^12^. The same analysis was not extended to the UCE dataset, which would have provided an informative control. Interestingly, our GGI analysis of TOGA2 single-copy exons, which is conceptually similar to ΔGLS in evaluating locus-by-locus support across predefined topological alternatives, favors FM overall. This shows that exonic data do not inevitably bias inference toward FP, even when analyzed under homogeneous-model frameworks. Second, Lü et al.’s syntenic block analysis used as a reference a *Psettodes* genome comprising 970 contigs with an N50 of 2.5 Mb, for chromosome-scale comparisons. Genome discontiguity is now recognized as a major driver of error in synteny-based inference^38^, and such fragmentation would be expected to inflate noise in the comparisons used to argue for polyphyly. This limitation motivated our generation of a chromosome-level genome for *Psettodes* erumei. Third, the higher syntenic similarity between *Psettodes* and non-flatfish carangarians reported by Lü et al. is fully consistent with FM. Under monophyly, *Psettodes* may retain the ancestral carangarian chromosome architecture, while some pleuronectoid taxa (most notably, *Solea*) underwent extensive derived rearrangement. This would produce exactly the pattern: greater syntenic conservation between *Psettodes* and outgroups than between *Psettodes* and pleuronectoids. Shared retention of the ancestral state should therefore not be conflated with sister-group relationship. Finally, Lü et al.’s syntenic comparison was distance-based, sidestepping the model-based phylogenetic framework that the field has developed precisely to handle the complexities of genomic data.

### Microsynteny as an independent phylogenomic data class

Gene-order rearrangements provide a class of phylogenomic evidence that is mechanistically independent of nucleotide substitution. Because the re-creation of an identical gene adjacency by random rearrangement is expected to be far less common than disruption of an existing adjacency, microsynteny characters carry low homoplasy^26^ and are less susceptible to the compositional artifacts that have confounded sequence-based flatfish phylogenomics. This property has been exploited to resolve other recalcitrant nodes: conserved gene linkage resolved the Ctenophora-versus-Porifera question^20^, chromosome-level rearrangements clarified the deepest teleost divergence^22^, and microsynteny networks reconstructed angiosperm phylogeny from over 120 genomes^25^.

In our analyses, microsynteny resolved the placement of *Psettodes* more decisively than it resolved the broader carangarian backbone. This patterns is consistent with the expectation that local gene-order rearrangements accumulate on a timescale relevant to the rapid succession of speciation events surrounding the origin of flatfishes. The syntenet ML tree, the AU test on the full microsynteny matrix, the Dollo-likelihood analysis, and the majority of AGORA metrics all converged on FM. This agreement from a data class orthogonal to nucleotide sequence provides a qualitatively distinct line of support for monophyly, one that cannot be attributed to substitution-model misspecification or compositional bias.

### Macrosynteny and the limits of ancestral chromosome reconstruction in rapid radiations

Macrosynteny (chromosome-scale rearrangements) provided an FM-leaning but inconclusive signal. Several factors contribute to this limited resolution. First, chromosome fusions and fissions are rare events, and the Carangaria radiation involved short internodes over which few such events are expected to have occurred^9,13,39^. Second, the most extensive chromosome-scale rearrangement in the dataset is concentrated in Pleuronectoidei, whose genomes rank among the most reshuffled vertebrate genomes^32^. This pattern resembles other lineage-concentrated bursts of chromosomal rearrangement against otherwise deeply conserved syntenic backgrounds, including in clitellate annelids (earthworms and leeches)^40^. This derived noise dilutes any synapomorphic signal that might unite *Psettodes* with Pleuronectoidei. Third, DESCHRAMBLER reconstructions are inherently reference-dependent, and no single extant genome can serve as a common reference for all three candidate topologies, necessitating the pairwise comparison design we adopted. Despite these limitations, adjacency log-likelihood favored FM in all four DESCHRAMBLER comparisons, and both *Psettodes*-reference comparisons (full-taxon and same-depth rerun) reached formal statistical support (paired Wilcoxon p = 0.007 and p = 0.008, respectively; RACF-cluster bootstrap 95% CI excluded zero in both cases).

### Concluding remarks

The three scales of genomic evidence examined here (nucleotide sequence, microsynteny, and macrosynteny) are concordant in the direction of FM, with hierarchically decreasing statistical resolution that mirrors the hierarchically increasing rarity of the underlying mutational events. Nucleotide substitutions are frequent and, when evaluated at the genome scale, provide strong support from the sheer volume of independent sites sampled genome wide. Gene-order rearrangements are rarer but carry low homoplasy and independently corroborate FM. Chromosome-scale fusions and fissions are rarest of all and, although FM-leaning, accumulate too slowly to resolve the short carangarian internodes decisively.

This hierarchical concordance pattern is interpretable in light of Carangaria’s evolutionary history. The clade represents a post-Cretaceous adaptive radiation along the benthic-pelagic axis^39,41,42^, in which the extinction of competitors following the K/Pg boundary opened ecological space that was rapidly partitioned among lineages as morphologically disparate as marlins, remoras, archerfishes, and flatfishes. Of the ∼1,100 carangarian species, flatfishes (>800 spp.) are the phenotypically most extreme example of this radiation. The short internodes produced by this rapid diversification provide sufficient time for microsyntenic rearrangements (local inversions, transpositions, and block movements) to accumulate as phylogenetic synapomorphies, but insufficient time for the rarer chromosome-scale fusions and fissions that macrosyntenic analyses require.

Together with the fossil record of gradual orbital migration^3^, multiple anatomical synapomorphies uniting all flatfishes^16–18^, and shared positively selected genes in two developmental pathways (the thyroid hormone pathway via TTF1 and BMP signaling via *bmp4*) that are detected only under the monophyly framework^12^, our multi-scale genomic evidence supports a single evolutionary origin of body plan asymmetry in flatfishes. Future work should prioritize generating additional VGP-quality chromosome-level assemblies for carangarians. It should also focus on developing and applying new tools for synteny-based phylogenomics, a field still in its infancy. In our analyses, individual tree-generating methods yielded locally weakly supported or implausible nodes even when the aggregate signal converged on the FM topology. These advances will improve our ability to reconstruct synteny conservation and transformation at any phylogenetic depth.

## Methods

### Genome assembly, annotation, and data acquisition

#### De novo genome assembly of *Psettodes erumei*

We produced a chromosome-level reference genome for *Psettodes erumei* (Indian spiny turbot), the only extant genus of Psettodoidei and therefore the key taxon for evaluating phylogenetic placements bearing on whether flatfish cranial asymmetry is more compatible with a single origin or multiple origins. Two specimens (one sinistral, KUMF 7742.1, 333.00 mm SL and one dextral, KUMF 7742.2, 237.00 mm SL) were collected at Ko Lek, Krabi Province, Thailand (Andaman Sea; 7.8128°N, 98.993°E) by P. Kantha in September 2024 at 6 m depth using a crab gill net. Tissue samples (muscle, gills, eye, and liver) were flash-frozen in liquid nitrogen at the collection site. Frozen tissues were stored at −80°C at the Department of Aquaculture, Kasetsart University, Bangkok, and subsequently shipped on dry ice to the UC San Diego (UCSD) Core Genomics Facility for DNA extraction, library preparation, and sequencing. Voucher specimens are deposited at KUMF (Kasetsart University Museum of Fisheries). Genomic DNA and all sequencing libraries used for the de novo assembly were derived from the dextral specimen.

Genomic DNA was extracted from muscle tissue using the NanoBind PanDNA kit (PacBio). Pacific Biosciences (PacBio) HiFi long-read sequencing was performed on a Revio system at the Stem Cell Genomics Core (gCore), Sanford Stem Cell Institute, UCSD, yielding approximately 116x coverage. Arima Genomics Hi-C proximity-ligation libraries were prepared following the Arima High Coverage Hi-C protocol and sequenced on a NovaSeq X 25B at the IGM Genomics Center, UCSD, yielding approximately 208x coverage (chromosome contact map in Supplementary Fig. S2). To aid genome assembly, Oxford Nanopore Technologies (ONT) reads for a scaffold-level genome of *P. erumei* were obtained from the NCBI Sequence Read Archive (SRR10913243; BioProject PRJNA592798), originally generated by the Center for Ecological and Environmental Sciences on a GridION platform (26.7 M reads, 47.1 Gb). PacBio HiFi reads, ONT reads, and Hi-C proximity-ligation data were jointly assembled with hifiasm v0.19.8^43^ using the -h1/-h2 flags to integrate Hi-C reads for haplotype phasing, leveraging the complementary long-read technologies to improve resolution of repetitive genomic regions. The assembly produced two phased haplotype contig sets (assembly statistics in Supplementary Table S11). Prior to scaffolding, each haplotype assembly was screened for contaminant sequences using Kraken2 v2.1.2^44^ against the standard Kraken2 database (archaea, bacteria, human, plasmid, UniVec_Core, and viral libraries), and identified non-target contigs were removed. Hi-C reads were then mapped to each cleaned haplotype assembly using the Arima Genomics mapping pipeline to generate the input alignments required for scaffolding. The cleaned and mapped haplotype assemblies were then scaffolded into chromosome-scale pseudomolecules with YaHS v1.2.2^45^.

Assembly completeness was assessed with BUSCO v6.0.0^46^ against the Actinopterygii odb10 database (Supplementary Table S12), yielding 98.5% complete BUSCOs (97.1% single-copy, 1.4% duplicated), 0.2% fragmented, and 1.3% missing. Base-level accuracy was evaluated with Merqury v1.3^47^ (k-mer distribution in Supplementary Fig. S1), producing a consensus quality value (QV) of 58.4. Contiguity statistics were computed with gfastats v1.3.6^48^: the final assembly comprised 24 chromosome-scale scaffolds totaling 588 Mb (98.6% of the total assembly length), with a contig N50 of 12.9 Mb (L50 = 31) and a scaffold N50 of 21.1 Mb (L50 = 13). Manual curation was performed in PretextView v0.2.5^49^ using Hi-C contact maps to identify and correct mis-joins and mis-scaffolds. The curated assembly was further screened for contamination using the NCBI Foreign Contamination Screen (FCS-GX)^50^ to remove any remaining non-target sequences prior to downstream analyses. After curation, hap2 was retained as the primary assembly on the basis of higher contiguity and completeness relative to hap1, and it was therefore used for repeat masking, annotation, whole-genome alignment, and accessioning; hap1 was retained only as a secondary phased representation. The final genome assembly and raw sequencing reads are available in NCBI (BioProject, assembly, and SRA accession numbers XXX, XXX, and XXX, respectively).

#### Additional genome assemblies

To broaden the phylogenomic framework within Carangaria, we incorporated two additional chromosome-level genomes: *Coryphaena hippurus* (mahi-mahi; 2n = 48; n = 24) and *Seriola dorsalis* (yellowtail amberjack; 2n = 48; n = 24). Both species were collected using rod and reel on a kelp paddy on August 22^nd^ 2021 by recreational anglers off of La Jolla CA 32°55’06.85”N 117°30’11.25”W. Blood samples were kept in 6 ml vacutainer K2 EDTA blood collection tubes (GTIN: 30382903678632, Becton Dickson) and transported back to the lab where they were held at 4C until extraction for a few days until they were extracted to maximize DNA yield and length^51^. DNA was extracted from 10 ul of whole blood within 3 days of collection using the NEB Monarch HMW kit for cells & blood (Cat# T3050L, NEB). Genomes for both species were sequenced using Oxford Nanopore Technologies (ONT) using the Ultra Long DNA sequencing kit (SQK-ULK001) and R9.4.1 chemistry on a PromethION 24 (minKnow version 21.05.12). This included two separate runs (L000-082, S108; L000-083 S105) for *S. dorsalis* and one run for *C. hippurus* (L000-081, S109). Standard Illumina libraries were generated using the Collibri NGS kit (Cat# A38602024, ThermoFisher) for *S. dorsalis* (L000-356) and *C. hippurus* (L000-355) (R000-080). Hi-C libraries were prepared for both *S. dorsalis* and *C. hippurus* (L000-426) (L000-355, L000-425) (R000-068).

Draft assemblies were constructed with Flye v2.9^52^, polished in three successive rounds of Racon^53^ followed by three rounds of Pilon^54^ using Illumina short reads, and scaffolded into chromosome-scale pseudomolecules with Hi-C proximity-ligation data. Assembly completeness was assessed with BUSCO v6.0.0^46^ against the Actinopterygii odb12 database, yielding 98.3% complete BUSCOs for *C. hippurus* and 97.8% for *S. dorsalis*. The *C. hippurus* assembly comprised 24 chromosome-scale scaffolds totalling 564 Mb, with a contig N50 of 18.4 Mb (L50 = 13) and a scaffold N50 of 24.3 Mb (L50 = 10); the *S. dorsalis* assembly comprised 24 chromosome-scale scaffolds totalling 653 Mb, with a contig N50 of 15.3 Mb (L50 = 16) and a scaffold N50 of 28.3 Mb (L50 = 11).

#### Taxon sampling and dataset assembly

A further 14 chromosome-level genome assemblies representing flatfishes and other carangarian lineages (full species and metadata list in Supplementary Tables S13, S14; specimen voucher and sequencing metadata in Supplementary Table S13) were retrieved from NCBI: *Scophthalmus maximus* (GCF_022379125.1), *Paralichthys olivaceus* (GCF_024713975.1), *Limanda limanda* (GCF_963576545.1), *Solea solea* (GCF_958295425.1), *Verasper variegatus* (GCA_026259375.1), *Eleutheronema tetradactylum* (GCA_046255195.2), *Lates calcarifer* (GCF_001640805.2), *Toxotes jaculatrix* (GCF_017976425.1), *Xiphias gladius* (GCF_016859285.1), *Rachycentron canadum* (GCA_038496675.1), *Echeneis naucrates* (GCF_900963305.1), *Trachinotus ovatus* (GCA_022709315.2), and *Trachurus trachurus* (GCA_905171665.2), along with *Mastacembelus armatus* (GCF_900324485.3; zig-zag eel; order Synbranchiformes) as a phylogenetic outgroup from Anabantaria, the sister group to Carangaria^4–9^. Together with the three newly assembled genomes, the final dataset comprised 17 species: six flatfishes (Pleuronectiformes: *P. erumei*, *S. maximus*, *P. olivaceus*, *L. limanda*, *S. solea*, and *V. variegatus*), ten non-flatfish carangarians (*E. tetradactylum*, *L. calcarifer*, *T. jaculatrix*, *X. gladius*, *R. canadum*, *C. hippurus*, *E. naucrates*, *T. ovatus*, *T. trachurus*, and *S. dorsalis*), and one outgroup (*M. armatus*). Genome sizes ranged from 538 Mb (*S. maximus*) to 801 Mb (*T. trachurus*). Haploid chromosome numbers ranged from n = 21 (*S. solea*) to n = 26 (*E. tetradactylum*), with the majority of species possessing n = 24.

#### Repeat masking and annotation

To prepare the three newly assembled genomes for gene annotation and whole-genome alignment with Progressive Cactus^36^, repetitive elements were identified de novo with RepeatModeler v2.0.5^55^ and the resulting species-specific repeat libraries were used to soft-mask each genome with RepeatMasker v4.1.7^56^ (repeat composition summary in Supplementary Table S15). Tandem repeats were additionally identified and masked with TRF v4.09.1^57^ using parameters 2 7 7 80 10 50 500 (match weight, mismatch penalty, indel penalty, match probability, indel probability, minimum score, maximum period); TRF output was parsed and used to soft-mask each genome. The resulting total repeat content was 30.2% (*P. erumei*), 13.1% (*C. hippurus*), and 23.2% (*S. dorsalis*). Total interspersed repeats accounted for 26.5% of the *P. erumei* genome, of which unclassified elements were the largest fraction (21.1%), followed by LTR retrotransposons (1.8%), DNA transposons (1.8%), LINEs (1.7%), and SINEs (0.1%). In *C. hippurus*, interspersed repeats comprised 9.1% of the genome, dominated by unclassified elements (4.6%), DNA transposons (1.9%), LINEs (1.4%), LTR retrotransposons (1.0%), and SINEs (0.3%). In *S. dorsalis*, interspersed repeats comprised 19.2%, with unclassified elements again the largest class (12.0%), followed by DNA transposons (4.2%), LINEs (1.4%), LTR retrotransposons (1.4%), and SINEs (0.2%).

#### Gene annotation

To annotate protein-coding genes in the three newly assembled genomes (*P. erumei*, *C. hippurus*, and *S. dorsalis*) and the five retrieved species lacking RefSeq-source annotations (*E. tetradactylum*, *V. variegatus*, *T. trachurus*, *T. ovatus*, and *R. canadum*), we employed two complementary ab initio and evidence-based pipelines: BRAKER3 v3.0.8^58^, which combines GeneMark-ETP and AUGUSTUS, and GALBA v1.0.11^59^, which uses 25iniport-based protein-to-genome alignments to guide AUGUSTUS predictions. No RNA-seq data were incorporated; both pipelines were run using protein evidence only. The protein evidence database was assembled by concatenating (i) the BUSCO v5 Actinopterygii odb10, (ii) Actinopterygii proteins extracted from OrthoDB v11, (iii) a curated panel of UniProt reference proteomes, and (iv) a locally collated set of fish proteomes from NCBI. Preliminary gene sets from each pipeline were merged and reconciled with TSEBRA v1.1.2.5^60^ using default configuration weights with the – filter_single_exon_genes and –ignore_tx_phase flags to produce a consensus annotation. TOGA2 (see below) became available after these annotations were completed and was therefore not incorporated as a third annotation source.

This procedure yielded 33,392 (*P. erumei*), 35,300 (*C. hippurus*), 39,965 (*S. dorsalis*), 32,759 (*E. tetradactylum*), 35,168 (*V. variegatus*), 36,211 (*T. trachurus*), 37,494 (*T. ovatus*), and 34,946 (*R. canadum*) protein-coding gene models for the eight genomes that lacked suitable RefSeq-source annotations. The same workflow was then applied to the nine remaining retrieved assemblies that already had RefSeq annotations, producing a fully harmonized 17-species annotation set for the final protein-based synteny analyses. Protein counts and BUSCO completeness scores for this annotation set are reported in Supplementary Table S16 (with assembly BUSCO in Supplementary Tables S11 and S12).

### Nucleotide-based, whole-genome phylogenomic analyses

#### Species tree inference with ROADIES

To obtain an initial species tree topology without requiring prior genome annotation or a reference genome, we applied ROADIES v1.0.0^28^ (Supplementary Fig. S3), a reference-free and alignment-free phylogenomic pipeline. Rather than relying on predefined markers or orthology detection, ROADIES identifies homologous regions directly from whole-genome assemblies by performing pairwise LASTZ^61^ alignments of randomly sampled genomic segments against each genome in the dataset. After quality filtering of the resulting alignments, ROADIES performs multiple sequence alignment with PASTA^62^, estimates both single-copy and multi-copy gene trees, and infers a species tree using ASTRAL-Pro2^63^, which explicitly accounts for both incomplete lineage sorting (ILS) and gene duplication or loss under a multi-species coalescent framework. ROADIES was run in deep mode starting from 2,000 genes for two iterations, with default parameters except for the minimum LASTZ alignment identity, which was increased to 85% (default 65%) to improve the proportion of high-confidence loci recovered.

#### Whole-genome alignment with Progressive Cactus

To evaluate the sensitivity of downstream phylogenomic inference to the assumed guide tree topology, we performed two independent whole-genome alignments using Progressive Cactus v2.9.7 (workflow and guide/reference configurations summarized in Supplementary Table S17)^36^. Progressive Cactus is a reference-free, multiple whole-genome aligner that constructs a hierarchical alignment graph without requiring a single reference genome; however, the algorithm uses a guide phylogeny to define the progressive decomposition of the alignment problem. The first alignment used the ROADIES-inferred species tree as guide, in which *P. erumei* is placed as sister to all remaining flatfishes (the monophyly hypothesis). For the second alignment, we manually modified the guide tree by repositioning *P. erumei* as sister to *T. jaculatrix* (a non-flatfish carangarian), reflecting the alternative topology reported by Lü et al.^15^, in which P. erumei is placed outside Pleuronectoidei.

Both alignments were performed on the repeat-soft-masked genomes with default Progressive Cactus parameters. The resulting Hierarchical Alignment (HAL) outputs were projected into multiple alignment format (MAF) using the hal2maf utility^64^. Because both the guide tree topology and the choice of reference genome for coordinate projection can introduce systematic biases into the extracted alignment blocks, we adopted a factorial design to disentangle these effects. For each HAL alignment, MAF alignments were extracted using two different reference species: (i) the outgroup *M. armatus*, which provides an outgroup-based reference projection, and (ii) *P. erumei* itself, the taxon of primary phylogenetic interest. This yielded four raw MAF alignments (two guide trees by two reference projections).

#### Assessment of guide-tree and reference bias

To perform an initial assessment of phylogenomic signal in the whole-genome alignments, we first converted the four raw MAF alignments to PHYLIP format with a custom script, then ran CASTER v1.0^27^ directly on them. The species trees inferred from these raw alignments consistently retained the local placement implied by the Progressive Cactus guide tree (Supplementary Figs. S5–S6) rather than an independently recovered phylogenetic signal. All monophyly-guided raw alignments resolved a topology in which *P. erumei* was placed as sister to the remaining flatfishes (Pleuronectoidei). In contrast, all polyphyly-guided raw alignments resolved topologies in which *P. erumei* fell outside the sampled flatfish clade, but they did not always reproduce the modified guide tree exactly: the raw CASTER-site analysis of the *P. erumei*-projected alignment placed *P. erumei* as sister to *T. jaculatrix*, matching the alternative guide tree, whereas the other polyphyly-guided raw analyses instead placed *P. erumei* as sister to the (*T. jaculatrix* + *X. gladius*) clade. This systematic correspondence between guide tree and inferred species tree indicated that guide-tree bias inherent to the Progressive Cactus alignment was propagating into downstream species tree inference. Reference-species bias was also detected: under the alternative guide tree, changing the MAF projection reference from *M. armatus* to *P. erumei* altered the raw CASTER-site result, whereas raw CASTER-pair remained polyphyletic under both reference projections. To mitigate these biases, we developed a post-processing pipeline to realign and filter the raw Progressive Cactus output before re-running CASTER.

#### Alignment post-processing

To mitigate residual guide-tree bias in the raw Progressive Cactus output, each MAF alignment was first filtered with a custom script to remove alignment blocks with more than 50% gaps, fewer than 50% of species represented, or fewer than 10 columns, then split into non-overlapping chunks of 25 kb, and converted to FASTA format. Each chunk was individually realigned with MAFFT v7.520^65^ using the L-INS-I strategy (--localpair –maxiterate 1000 –ep 0.123 – anysymbol) to reduce residual guide-topology dependence during local homology refinement, and trimmed with trimAl v1.4.1^66^ using the -nogaps flag to retain only gap-free columns. After trimming, only sites with complete taxon occupancy across all 17 species were retained. All downstream post-processed CASTER analyses were then conducted on the *P. erumei*-projected alignments. To test whether the final species-tree estimates still retained topological dependence on the original Cactus guide tree, we compared the resulting post-processed CASTER trees to both guide trees and to one another using unrooted Robinson-Foulds (RF) distances (Supplementary Figures S9 and S10). These RF comparisons showed that the post-processed trees did not simply recapitulate their input guide trees: both post-processed CASTER-site trees were identical (RF = 0), and all four final CASTER trees were closer to the monophyly-guided tree (RF = 2-4) than to the alternative-guided tree (RF = 6-10). Together, these comparisons indicate substantial convergence after realignment and trimming and suggest reduced guide-topology dependence in the post-processed alignments. The final concatenated alignments comprised 122,287,546 sites (monophyly-guided) and 121,949,235 sites (polyphyly-guided), of which 33,065,715 and 32,967,064 were parsimony-informative, respectively.

#### Coalescent-based species tree estimation with CASTER

Following alignment post-processing, we re-ran coalescent-based species tree inference on the realigned data using CASTER v1.0^27^. Unlike conventional coalescent methods that first estimate individual gene trees and then reconcile them into a species tree, CASTER estimates the species tree directly from whole-genome alignment site patterns. We applied two complementary CASTER algorithms: CASTER-site, which estimates the species tree from patterns of unlinked biallelic sites across the genome, and CASTER-pair, which estimates the species tree from pairwise coalescence time estimates derived from the alignment. Both methods were applied independently to each of the two post-processed alignments (monophyly-guided and polyphyly-guided), yielding four independent species tree estimates in total. The CASTER-pair analysis of the polyphyly-guided alignment yielded an unexpected topology in which *E. tetradactylum* was placed as sister to Pleuronectoidei; this *Eleutheronema* + Pleuronectoidei placement was not a pre-specified hypothesis and arose as a post hoc result from this single analysis, and was subsequently carried forward as a third alternative topology (FP_1_) in the comparative analyses described below.

#### Single-copy ortholog identification with TOGA2 and gene-tree/species-tree concordance

To obtain a high-confidence set of single-copy orthologs for independent phylogenomic validation and as input for ancestral gene-order reconstruction, we generated pairwise whole-genome alignments in UCSC chain format for each of the 17 species against the three-spined stickleback (*Gasterosteus aculeatus,* GCF_964276395.1) using make_lastz_chains (Kirilenko et al., 2023) with LASTZ v1.04.52^61^ and default chaining parameters. The stickleback was chosen as the projection reference because of its well-annotated, chromosome-level genome and its phylogenetic position outside Carangaria, minimizing the potential for reference bias within Carangaria. Chain-format alignments were used as input for TOGA2 v2.0.7^29^, which projects reference gene annotations onto query genomes through whole-genome alignment to classify genes as one-to-one orthologs, many-to-one orthologs, or paralogs, and assigns each classification a confidence score based on alignment coverage, synteny context, and sequence identity. From the TOGA2 results, we retained only one-to-one orthologs with an orthology confidence score ≥ 0.99 (the highest confidence tier, indicating genes that are intact, non-duplicated, and well-supported by synteny) that were present in all 17 species, yielding 10,430 high-confidence single-copy ortholog families.

For each ortholog family, coding sequences were extracted and aligned with MAFFT v7.520^65^ using the –auto strategy. Aligned sequences were trimmed with trimAl v1.4.1^66^ using the -automated1 heuristic. We used the trimmed coding-sequence alignments to infer a concatenation-based maximum-likelihood phylogeny. Alignments were concatenated across the 10,430 single-copy orthologs, and the supermatrix was analyzed in IQ-TREE 2^67^ under a partition scheme in which each gene was partitioned by codon position. Branch support was assessed with 1,000 ultrafast bootstrap replicates. Individual gene trees were inferred for each of the 10,430 orthologs with IQ-TREE 2.2.2.7 (full ASTRAL concordance-factor tree in Supplementary Fig. S11; key-node support in Supplementary Table S18)^67^ under the best-fit substitution model selected by ModelFinder^68^ using the Bayesian Information Criterion (BIC), with 1,000 ultrafast bootstrap replicates (-B 1000) and 1,000 SH-aLRT replicates (-alrt 1000) to assess branch support. A species tree was then reconstructed from the full set of 10,430 gene trees using the multi-species coalescent implemented in ASTRAL-III v5.7.1^69^. To quantify the robustness of the focal relationship, we calculated gene concordance factors (gCF) and site concordance factors (sCF) using the –gcf and –scf options in IQ-TREE 2^67^.

#### Gene genealogy interrogation of competing topologies

To evaluate relative support for each of the three competing topologies across individual loci, we applied an approach inspired by gene genealogy interrogation (GGI;^30^) to the 10,430 single-copy ortholog alignments. Rather than exhaustively enumerating all possible trees, we adopted a targeted variant that restricted the evaluation to the three topological hypotheses motivated by the flatfish-origins debate (per-gene AU-ranking curves in Supplementary Fig. S12; Supplementary Table S19). We evaluated three constraint topologies that each fixed a single disputed node while leaving the rest of the tree free to be optimized: (i) *P. erumei* constrained as sister to Pleuronectoidei, consistent with FM; (ii) *P. erumei* constrained as sister to *T. jaculatrix* (FP_2_); and (iii) *E. tetradactylum* constrained as sister to Pleuronectoidei (FP_1_). For each of the 10,430 loci, we inferred the best tree under each constraint with IQ-TREE 2 (Minh et al., 2020) using the best-fit substitution model identified by ModelFinder^68^, and compared the constrained tree to the unconstrained maximum-likelihood tree using the approximately unbiased (AU) test with 10,000 RELL bootstrap replicates (-zb 10000 -au). A constraint was considered rejected for a given locus when the AU test p-value fell below 0.05.

### Synteny-based analyses

#### Data preparation

Before finalizing the protein-based synteny datasets, we first tested whether annotation provenance distorted downstream microsynteny-based phylogenetic inference. We initially applied the microsynteny workflow described below to a mixed annotation set that combined RefSeq models for species with available NCBI gene models and our newly generated annotations for the remaining taxa. Because that exploratory analysis grouped species by annotation source rather than by phylogenetic affinity (Supplementary Fig. S13), we repeated the same test with a second mixed regime in which some species were represented only by BRAKER annotations and others only by GALBA annotations (Supplementary Fig. S14; species assignments in Supplementary Table S19-S20). The recurrence of source-driven clustering in both cases indicated that heterogeneity in gene boundaries, exon structures, and isoform definitions was distorting protein-based synteny inference. We therefore standardized annotation source before conducting the final microsynteny and macrosynteny analyses.

The nine assemblies that already carried RefSeq annotations were re-annotated with the same unified pipeline described in the annotation section, yielding a fully harmonized 17-species proteome set. All assemblies were restricted to chromosome-scale scaffolds, each gene was represented by a single isoform, and simplified annotation tables recording scaffold, strand, and genomic coordinates were generated for downstream use. The same harmonized dataset was then used for the gene-order and chromosome-scale protein-based synteny analyses described below. The four data streams described in this section feed into two downstream analytical scales: the harmonized proteomes and the TOGA2-derived ortholog tables were used for the microsynteny analyses, whereas the reciprocal-best-hit ortholog sets and the whole-genome alignment chains were used for the macrosynteny analyses. Protein counts and BUSCO completeness scores for this harmonized set are reported in Supplementary Table S21.

For the gene-order analyses, we additionally prepared ortholog gene-order tables from the TOGA2 projections^29^. A custom script extracted one-to-one orthologs classified as Fully Intact (FI), Intact (I), or Partially Intact (PI) and reformatted them into orthology groups and extant gene-order tables. We prepared a strict set retaining only orthologs with score ≥ 0.99 and a relaxed set with no score filter, as described in the corresponding analysis subsections. For the pairwise chromosome-scale comparisons, we generated reciprocal-best-hit ortholog sets from the same harmonized proteomes. For each species pair, all-versus-all DIAMOND v2.1.13 BLASTP^70^ searches were carried out and only reciprocal-best-hit orthologs were retained, selecting the top-scoring match in each direction by bit score and e-value. For the alignment-based ancestral chromosome reconstruction, pairwise whole-genome alignment chains were exported from the Progressive Cactus HAL files^36,64^ in UCSC chain format for each reference genome using cactus-hal2chains. Chromosome-size tables were exported with halStats^64^. Chain sets were reoriented when necessary with chainSwap and then processed through chainMergeSort, chainPreNet, chainNet, and netSyntenic^71^, producing the per-chromosome chain/net hierarchy used in that downstream analysis.

#### Microsynteny analyses

For microsynteny analyses, we used two independent analytical frameworks: syntenet, which clusters protein-based collinear blocks into a binary microsynteny character matrix, and AGORA, which reconstructs gene-order relationships from TOGA2-derived single-copy orthologs.

##### Syntenet

###### Microsynteny network phylogeny (tree-generating)

All-versus-all DIAMOND v2.1.13 BLASTP searches^70^ were conducted across all 17 filtered proteomes without reciprocal-best-hit filtering, allowing a broader set of homologous matches to contribute to local collinearity detection. A custom shell script filtered each similarity table to retain only hits with amino-acid identity > 70% and aligned length > 30 amino acids. The resulting filtered similarity tables together with the harmonized coordinate tables were then supplied to MCScanX v1.0.0^72^, retaining up to five best non-self hits per query, requiring at least five anchors per collinear block, allowing a maximum gap of 25 genes, and collapsing overlapping matches within a five-gene window; all other settings were left at defaults.

The resulting syntenic anchor pairs were then analyzed with the R package syntenet v1.3.0^31^ for network-based clustering across taxa. Genes connected by conserved syntenic relationships form an edge list that is clustered with the Infomap algorithm^73^ to define synteny clusters. These clusters were then converted to a binary presence/absence matrix in which rows corresponded to species and columns to synteny clusters. The final matrix contained 23,850 binary microsynteny characters across the 17 taxa and was exported in PHYLIP format for downstream phylogenetic analysis. We then inferred a phylogeny from the binary matrix with the infer_microsynteny_phylogeny function in syntenet^31^, which calls IQ-TREE 2 (Minh et al., 2020) under the Mk model^74^ for binary characters with free and equal rates and state frequencies (-m Mk+R+FO). Branch support was assessed with 1,000 ultrafast bootstrap replicates and 1,000 SH-aLRT replicates.

###### AU test on the microsynteny character matrix (topology-dependent)

The same full microsynteny character matrix was then used to compare the three topologies directly via the approximately unbiased (AU) test. We retained the full 23,850-character matrix and re-optimized each constrained topology in IQ-TREE under the same settings used for tree inference (--seqtype MORPH -m MK+FO+R) before evaluating the optimized candidate trees with 10,000 RELL replicates.

###### Cluster profile counts (topology-dependent)

To summarize occupancy patterns within the microsynteny cluster matrix, we quantified the clustered phylogenomic profile matrix and visualized a variant-only, capped version of it (Supplementary Figure S17). For this descriptive occupancy-based summary, we excluded clusters present in only a single taxon or in more than 90% of taxa, thereby reducing noise from lineage-specific singletons and nearly ubiquitous clusters. Of the 23,850 clusters in the binary matrix, 20,579 were retained after applying this filter, and topology-exclusive cluster counts under FM, FP_1_, and FP_2_ were tallied from this filtered set.

##### AGORA

The gene-order analyses used the TOGA2-derived ortholog gene-order tables, formatted for AGORA compatibility as described above, as their starting point. Running AGORA^32^ on these tables produced two distinct types of output: (i) extant gene adjacencies extracted at the tips of the tree, which describe the observed gene orders in the sampled genomes and do not depend on the assumed phylogeny, and (ii) ancestral gene-order reconstructions at internal nodes, which are conditioned on the input tree topology and were therefore computed separately for each of the three candidate topologies. Because previous AGORA-based vertebrate comparisons identified soles as exceptionally rearranged^32^, we retained *S. solea* in the primary analyses but repeated the gene-order summaries most sensitive to lineage-specific rearrangement in an *S. solea*-excluded sensitivity framework.

###### Rearrangement-distance minimum-evolution tree and binary adjacency maximum-likelihood tree (tree-generating)

Using extant tip adjacencies, we evaluated the strict TOGA FI/I/PI marker series at orthology-score cutoffs of 0.80, 0.95, and 0.99 to select a conservative ortholog set for downstream gene-order inference. At 0.80 (6,230 marker families; 9,155 adjacency characters), exploratory ML analysis of the binary adjacency matrix yielded a biologically implausible topology and did not recover any of the competing focal alternatives. At 0.95 (5,921 marker families; 8,485 adjacency characters), ML resolved FM with weak support (24%). At 0.99 (5,103 marker families; 7,023 adjacency characters), FM was again resolved with slightly higher support (32%). We therefore adopted the most stringent 0.99 cutoff for both the distance-based and character-based gene-order analyses to minimize the influence of uncertain orthology assignments.

In the full 0.99 dataset, we summarized adjacency structure as a taxon-by-taxon, normalized adjacency-distance matrix and inferred a balanced minimum-evolution tree with FastME^75^ from 1,000 bootstrap distance matrices. We next inferred an ML tree directly from a binary gene-adjacency matrix using the same 0.99 ortholog set. Using the relaxed vertebrates extant adjacency tables, we retained unsigned scaffold-internal adjacencies across all taxa and encoded 7,023 binary characters for analysis (Supplementary Figs. S24, S25; Supplementary Table S3) in IQ-TREE 2^67^ under –seqtype BIN with a requested JC2+R+FO model (fitted model: JC2+FQ+R4); support was assessed with 1,000 ultrafast bootstrap replicates and 1,000 SH-aLRT replicates. Constrained three-topology tests were then run on the same all-scaffolds adjacency matrix using IQ-TREE (--seqtype BIN -m JC2+R+FO), and topologies were compared with the AU test using 10,000 RELL replicates.

###### Ancestral gene-order reconstruction metrics (topology-dependent)

The second type of AGORA output is conditioned on the input tree topology. To reconstruct ancestral gene order under each candidate topology and evaluate how inferred ancestral chromosome segments varied around the disputed node, we ran AGORA^32^ on each candidate tree. AGORA infers ancestral gene adjacencies from conserved ordered orthologs across extant species and assembles them into Contiguous Ancestral Regions (CARs) (AGORA metric dashboard in Supplementary Tables S5, S4), that is, inferred ancestral chromosome segments rather than finished chromosomes. We evaluated four AGORA configurations to test sensitivity to ortholog filtering, workflow choice, and taxon sampling: (i) a strict vertebrates run retaining orthologs with score ≥ 0.99 (5,103 orthogroups), (ii) a relaxed generic run (11,110 orthogroups), (iii) a relaxed vertebrates run (11,110 orthogroups; our primary configuration), and (iv) an *S. solea*-excluded relaxed vertebrates sensitivity run (11,605 orthogroups).

For each competing topology, we computed CAR count and contiguity statistics (G50, L70); Jaccard similarity between the focal ancestor and the extant P. erumei adjacency set; incident breakpoints (adjacency disruptions on the branch entering the focal ancestor plus the path to extant P. erumei and the complement child) and surrounding branch breakpoints; and an extant adjacency parsimony analysis in which the same 11,190 informative adjacency characters were scored on each candidate topology and compared by total parsimony steps with 1,000 bootstrap replicates and a species jackknife. For each pair of focal reconstructions we also computed directed projection costs, the minimum number of destination CARs required to cover each large (≥ 20-gene) source CAR at 95% coverage. Focal-branch complexity was characterized by counting clear fissions, clear fusions, and ambiguous block-partition components on the branch entering each focal ancestor and on each descending branch.

#### Dollo likelihood analysis of gene adjacencies

To compare the three candidate topologies under a one-gain/multiple-loss model for adjacency evolution, we analyzed the relaxed vertebrates marker set in a Dollo-likelihood framework. This model is appropriate for exact gene-adjacency characters because the independent re-creation of the same ordered adjacency by rearrangement is expected to be rare relative to disruption of an existing adjacency, making repeated losses more plausible than repeated gains. We extracted a shared extant adjacency matrix from the relaxed vertebrates gene-order tables. The eight ingroup species used to define derived adjacencies were *L. limanda*, *P. olivaceus*, *S. maximus*, *S. solea*, *P. erumei*, *E. tetradactylum*, *T. jaculatrix*, and *L. calcarifer*, and the seven external background taxa used for outgroup filtering were *R. canadum*, *C. hippurus*, *E. naucrates*, *T. ovatus*, *T. trachurus*, *S. dorsalis*, and *M. armatus*. Across these taxa, we restricted the dataset to 17,280 shared marker families and 31,351 scaffold-internal adjacencies, then retained 5,922 derived adjacency characters present in at least one ingroup species but absent from all seven background taxa. This outgroup-based filtering enriches for adjacencies likely gained within the focal ingroup without privileging any of the three ingroup topologies.

Each candidate topology (FM, FP_1_, FP_2_) was evaluated on the 5,922-character adjacency matrix under a three-state, Dollo-type structured Mk model implemented in phytools::fitMk (R package phytools)^76^. States represent adjacency absent before gain (0), adjacency present (1), and adjacency absent after loss^77^. Transition rates were constrained to allow gains only from 0→1 and losses only from 1→2, with no regain after loss. Because all retained characters are absent from the background taxa by construction, the root state was fixed to 0 (pre-gain absence). Tip presences were coded as state 1, whereas tip absences were coded as ambiguous between states 0 and 2 (via a tip-likelihood matrix). For each candidate tree, gain and loss rates were optimized by maximum likelihood.

#### Macrosynteny analyses

For macrosynteny analyses (workflow in Supplementary Fig. S26), we used two complementary approaches: protein-based chromosome comparisons based on the harmonized proteomes and coordinate tables, and whole-genome-alignment-based ancestral chromosome reconstruction. The protein-based analyses were carried out in Oxford Dot Plots v0.3.7^20^, whereas the alignment-based ancestral reconstructions were performed with DESCHRAMBLER^33^.

##### Protein-based chromosome-scale comparisons

Protein-based chromosome-scale comparisons were conducted in Oxford Dot Plots v0.3.7^20^ using reciprocal-best-hit orthologs identified from the harmonized proteomes and chromosome coordinate tables described in Data preparation. This approach yielded two complementary lines of evidence: tree-independent summaries of extant chromosome correspondences, examined first through a direct multi-species comparison based on inferred ancestral linkage groups (ALGs) and then through pairwise comparisons; and topology-dependent tests of whether rearrangements defined within this ALG framework were consistent with any of the competing phylogenetic hypotheses.

###### Carangaria-specific ancestral linkage groups (descriptive)

To reconstruct this shallower reference system, we performed overlapping four-way analyses from the same harmonized proteomes and coordinate tables (anchor-assignment rules summarized in Supplementary Table S6), again retaining only reciprocal-best-hit ortholog groups. *L. calcarifer*, *P. erumei*, and *T. ovatus* were used as shared anchor taxa because they represent distinct carangarian families while all retaining the predominant n = 24 karyotype, thereby providing a taxonomically broad but chromosomally stable backbone for ALG inference. A fourth species was then rotated among *M. armatus*, *T. jaculatrix*, *T. trachurus*, *R. canadum*, *X. gladius*, *P. olivaceus*, *V. variegatus*, and *S. dorsalis*; taxa with derived haploid chromosome numbers other than n = 24 were excluded from this discovery set to minimize bias from lineage-specific chromosome-number change in defining the Carangaria-specific linkage system. Orthology searches were carried out with DIAMOND v2.1.13 BLASTP^70^, and only ortholog sets recovered as complete four-way reciprocal-best-hit groups were retained. These orthologs were grouped by the combination of scaffolds on which they occurred in the four species, producing candidate chromosome groups for each quartet. Significance of group size was assessed by shuffling gene IDs 1,000,000 times per quartet to estimate a group-size-specific FDR, and only groups whose size exceeded the 0.05 threshold were retained. ALG candidates were retained only when the same scaffold combination recurred across multiple rotating-taxon analyses, not on the basis of any single quartet. Across the eight analyses, these procedures converged on 24 Carangaria-specific ALGs numbered ALG01 through ALG24 according to *P. erumei* scaffold order.

###### Multi-species chromosome comparison (descriptive)

To trace these 24 ALGs across all 17 taxa, the quartet-derived reciprocal-best-hit ortholog groups were merged across analyses and extended to the remaining species with an HMM-based homology search. Each merged ortholog set was aligned with MAFFT^65^, converted into an HMM with HMMER v3.4^78^, and searched against the proteomes of the remaining species so that the best-scoring significant homolog could be recovered even in taxa not used in the quartet discovery phase. Missing data were permitted when no significant hit was recovered. This procedure produced a final expanded multi-species ALG homology table in which each ortholog set retained its curated Carangaria-specific ALG identity and, when recoverable, its gene, scaffold, and genomic position in every species of the dataset.

###### Pairwise chromosome-scale synteny conservation (descriptive)

To evaluate chromosome-scale structure across the dataset, we grouped reciprocal-best-hit orthologs for each species pair (Supplementary Table S10) by the combination of chromosome-scale scaffolds on which they occurred. Significance of group size was assessed by shuffling gene IDs in the corresponding .chrom coordinate tables 1,000,000 times per comparison to estimate a group-size-specific false discovery rate (FDR), and only groupings whose size exceeded the 0.05 FDR threshold were retained. These significant pairwise correspondences were then visualized as homologous scaffold pairs. Chromosomes were interpreted in light of the previously characterized Bilaterian-Cnidarian-Sponge linkage groups (BCnS LGs)^24^ and chordate linkage groups (CLGs)^23^, allowing us to distinguish deeply conserved linkage identities from lineage-specific derived combinations. Because the BCnS LG and CLG frameworks were too deep to resolve the more recent rearrangements relevant to the focal hypothesis tests, a shallower Carangaria-specific chromosome reference system was required for the downstream multi-species comparison.

###### Genome rearrangement simulations (topology-dependent)

Using the curated set of Carangaria-specific ALGs, we next asked whether any fusion-with-mixing events could be identified (quartet sims in Supplementary Figs. S29-31; quintet sims in Supplementary Figs. S32, S33) that were consistent with, and therefore potentially informative for, one of the three competing topologies. The rationale was that, if a derived chromosome combining segments from different ALGs arose on the branch supporting one of these alternative relationships, the descendant taxa united by that branch under the corresponding topology should share the resulting mixed-fusion signature. A fusion was scored as “mixed” when genes assigned to different ancestral linkage groups occupied overlapping coordinate ranges on the same derived chromosome. We then identified mixed fusions whose phylogenetic distribution was compatible with each candidate topology and scored each event by the total number of genes contributing to topology-compatible mixed fusions within a given taxon set. Simulations were carried out across multiple quartet and quintet configurations designed to evaluate the three alternative sister-group relationships represented by FM, FP_1_, and FP_2_. In all cases, *M. armatus* was used as the outgroup to polarize rearrangement states, at least one pleuronectoid species was included, and the remaining ingroup taxa were varied across alternative taxon sets so that the same topological predictions could be tested repeatedly under different sampling combinations. For each quartet or quintet test, we evaluated a null model in which protein IDs were shuffled among chromosomal coordinates within one genome at a time and fusion-with-mixing was rescored after each shuffle using the same overlap criterion. We performed 1,000,000 randomizations for each individual quartet and for each individual quintet. Empirical false discovery rates were then estimated as the proportion of shuffled trials that produced topology-support scores equal to or greater than the observed value for the corresponding quartet or quintet.

##### DESCHRAMBLER ancestral chromosome reconstruction

The whole-genome-alignment-based chromosome analyses used the per-chromosome chain/net hierarchies prepared from the Progressive Cactus HAL alignments (see Data preparation) as their starting point. Running DESCHRAMBLER^33^ on these inputs produces ancestral predicted chromosome fragments (APCFs) at internal nodes, but these reconstructions are conditioned both on the assumed topology and on the descendant reference genome used to anchor the coordinate system. This reference dependence meant that no single extant genome could serve as a common reference for all three candidate topologies, and it therefore defined the comparative design described below.

###### Reference-specific reconstruction design and resolution selection

Because no single extant reference could be used to reconstruct the focal ancestor under all three candidate topologies, we decomposed the three-topology problem into two reference-specific pairwise comparisons that share FM: a *Scophthalmus*-anchored FM versus FP_1_ comparison and a *Psettodes*-anchored FM versus FP_2_ comparison. *S. maximus* is a valid descendant reference for the focal ancestor under FM and FP_1_ but not under FP_2_, whereas *P. erumei* is a valid descendant reference under FM and FP_2_ but not under FP_1_. Among the five pleuronectoid descendants in the dataset, *S. maximus* was chosen as the FM/FP_1_ reference because it retains the most conserved chromosome-scale macrosynteny. Within each reference framework, ancestors were reconstructed at three resolutions (100 kb, 300 kb, and 500 kb). We selected 100 kb as the working resolution because, across all downstream runs, it consistently recovered the greatest fraction of the reference genome in the reconstructed ancestor (94.0% for 100 kb versus 88.5% for 300 kb and 84.8% for 500 kb); summary metrics for all reference-by-topology-by-resolution combinations are provided in Supplementary Table S7.

###### Comparison designs and evaluation metrics (topology-dependent)

Using the 100 kb reconstructions, we evaluated two DESCHRAMBLER comparison designs (evidence matrices in Supplementary Figs. S35–S38; Supplementary Tables S8–S23). The first used full taxon sampling. In the *Scophthalmus*-reference framework, this compared FM with FP_1_ using five pleuronectoid descendants in both runs. In the *Psettodes*-reference framework, it compared FM with FP_2_, but this comparison was asymmetric because the FM reconstruction used five descendants whereas the FP_2_ reconstruction used only *T. jaculatrix*. To account for this, we also repeated both reference frameworks in a two-descendant rerun, using the reference plus one focal descendant together with 11 outgroups, so that both within-reference comparisons were same-depth. For each topology within each design, we computed APCF count, reference coverage, APCF N50, counts of large and small fragments, fragmentation percentage, total SF count, and mean blocks per APCF, together with descendant-versus-outgroup adjacency-retention separation, breakpoint totals, switching-taxon diagnostics, adjacency log-likelihood, leave-one-out jackknife robustness, an APCF-cluster bootstrap, and a strict paired coordinate-matched adjacency test. Because APCF coordinates are reference dependent, strict paired tests were restricted to comparisons within each reference framework.

## Acknowledgements

Computational support for this project was provided by the University of Oklahoma Supercomputing Center for Education and Research (OSCER) and the Expanse system at the San Diego Supercomputer Center (SDSC). We thank Mark Miller (SDSC) for support and advice.

